# CRAK-Velo: Chromatin Accessibility Kinetics integration improves RNA Velocity estimation

**DOI:** 10.1101/2024.09.12.612736

**Authors:** Nour El Kazwini, Mingze Gao, Idris Kouadri Boudjelthia, Fangxin Cai, Yuanhua Huang, Guido Sanguinetti

**Affiliations:** Theoretical and Scientifc Data Science, Scuola Internazionale Superiore di Studi Avanzati, Trieste, Italy; School of Biomedical Sciences, The University of Hong Kong, HKSAR; Department of Statistics and Actuarial Science, The University of Hong Kong, HKSAR

**Keywords:** RNA velocity, chromatin accessibility, region kinetics

## Abstract

RNA velocity has recently emerged as a key tool in the analysis of single-cell transcriptomic data, yet connecting RNA velocity analyses to underlying regulatory processes has proved challenging. Here we propose CRAK-Velo, a semi-mechanistic model which integrates chromatin accessibility data in the estimation of RNA velocities. CRAK-Velo provides biologically consistent estimates of developmental flows and enables accurate cell-type deconvolution, while additionally shining light on regulatory processes at the level of interactions between genes and chromatin regions.

## 1 Main

Recent advances in single-cell sequencing technologies are leading to unprecedented insights in the molecular biology of cells [1–10]. However, the intrinsically destructive nature of sequencing creates a significant limitation in using such technologies to elucidate the dynamics of cellular processes. To circumvent this problem, La Manno et al. [11] proposed the concept of *RNA velocity*, a ground-breaking method which exploits splicing information to determine whether individual genes are being up-regulated or repressed at the time the sequencing happened. When aggregated at the cell level, this information provides valuable insight into the developmental or adaptation processes happening through the population of cells.

The RNA velocity concept has been extended and refined in a number of recent papers [12–17], yet all of these methods need to specify or learn from data the transcription rate of individual genes in each cell. This either imposes significant constraints on the models (when the transcription rate is postulated to be e.g. a binary switch), or greatly expands the number of parameters required by the model. Alternatively, one might use additional epigenomic information such as chromatin accessibility to explain transcription rates; such information is increasingly becoming available through multiomic single-cell techniques such as SNARE-seq[18], SHARE-seq[15], and the 10x Genomics Multiome commercial kit. This idea has been recently exploited by MultiVelo [19], an elegant model which aggregates chromatin accessibility and expression data through a differential equations latent variable model.

Here we present CRAK-Velo (ChRomatin Accessibility Kinetics integration in RNA Velocity), a simpler model which directly integrates chromatin accessibility data in the estimation of individual gene transcription rates. We draw inspiration from UniTVelo [12], a recently proposed RNA velocity model which estimates transcription rates using a parametric approximation, but expand it by explicitly connecting transcription rates to observed open region probabilities (quantified using a probabilistic topic model [20, 21]; see Methods for the mathematical formulation of the model). This achieves two main goals: it regularises estimation of transcription rates, ensuring it is consistent with chromatin information, and it quantifies the impact of each regulatory region during transcriptional dynamics.

Figure 1 shows the application of CRAK-Velo to a 10x Multiome data set of human stem cells undergoing hematopoietic differentiation [19]. Panel (a) shows the visualisations of the data set obtained by three different methods: CRAK-Velo (left), UniTVelo (centre) and MultiVelo (right). This challenging benchmark was used in [19] (cf Fig 5 in the original publication) to demonstrate how integrating chromatin accessibility could correct spurious velocity estimates. We also observe that the purely RNA-based method UniTVelo cannot correctly identify the platelet state as a terminally differentiated state in this data set. CRAK-Velo is however also able to identify three separate terminal states (granulocytes, erythrocytes and platelets), while MultiVelo detects a biologically dubious flow from erythrocytes to granulocytes. This and other observations can be quantified using a cross-boundary direction (CBDir) analysis, which compares inferred transitions with expected transitions from biological knowledge. Figure 1(b) shows histograms of this metric for the three methods, clearly showing the superiority of the flows inferred by CRAK-Velo compared to the other methods. We then considered the usefulness of the CRAK-Velo representation to distinguish cell types. Figure 1(c) shows the standard RNA velocity phase portraits (spliced vs unspliced read counts) for selected genes, while Fig 1(d) shows unspliced reads vs chromatin effect for the same genes for CRAK-Velo (top) and MultiVelo (bottom).

**Fig. 1.**
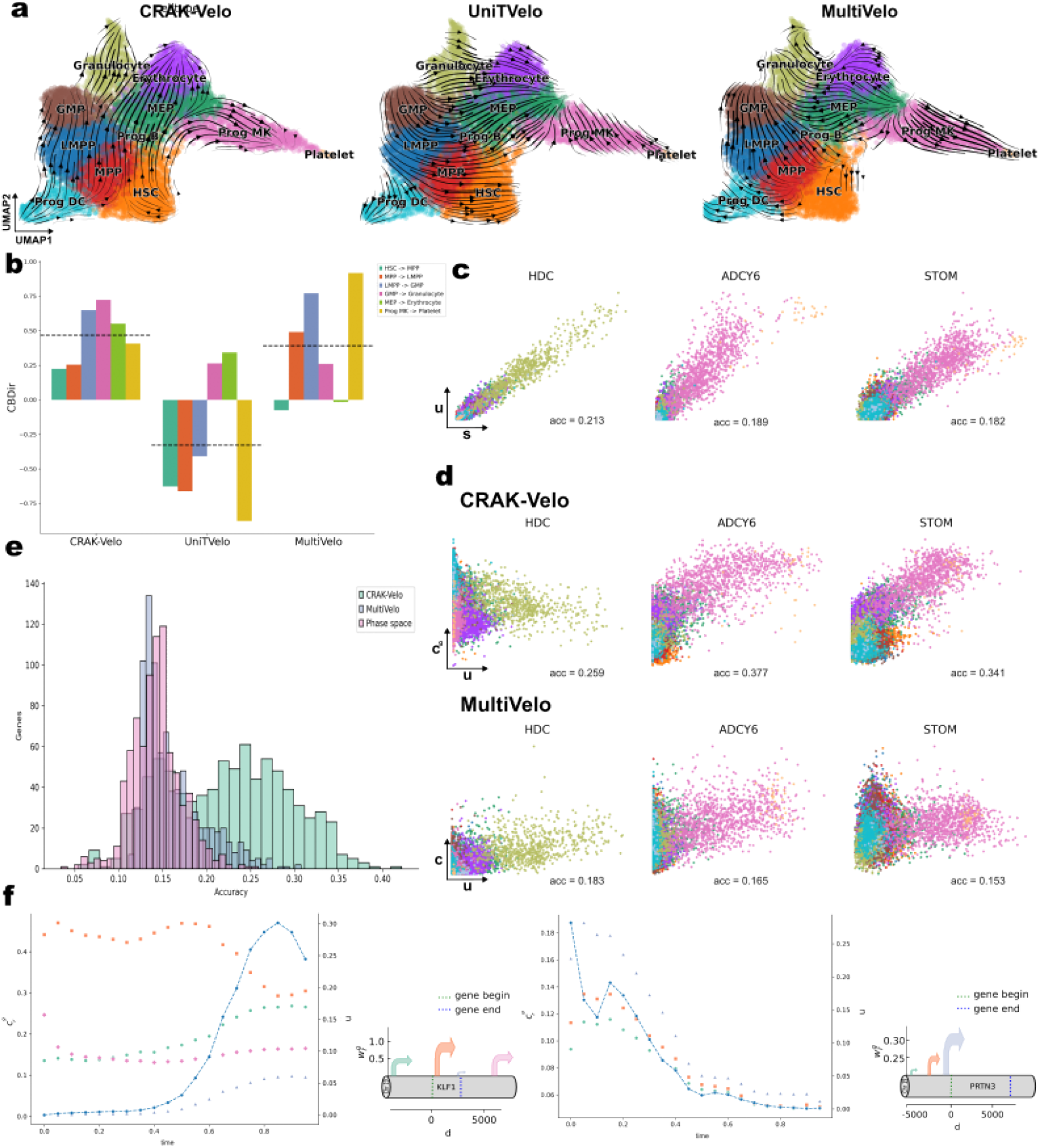
(a) Visualizations of velocity field showing inferred HSPC differentiation on the Umap embedding using three methods: CRAK-Velo (left), UniTVelo (center), and MultiVelo (right). (b) Histograms of cross-boundary direction (CBDir) analysis for the three methods, comparing inferred transitions with expected biological transitions. The black dashed line is the average CBDir per method. (c) Standard RNA velocity phase portraits (spliced vs. unspliced read counts) for selected genes. (d) Comparison of unspliced reads vs. chromatin effect for the same genes using CRAK-Velo (top) and MultiVelo (bottom). The number below each plot is the accuracy of a KNN cell type classifier. (e) Histogram of the performance of the KNN cell type classifier across all genes for the phase space and accessibility-unspliced space inferred by the CRAK-Velo and MultiVelo. (f) Dynamics of regulation for KLF1 and PRTN3 genes. Left is the region kinetic plot with double y axis that shows the average accessibility 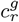 for each region around the gene on the left and the average normalised unspliced reads *u*_*g*_ on the right one. On the right is an arrow plot. The position of an arrow corresponds for start of a region and thickness is its contribution/weight 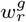 inferred by CRAK-Velo.

Different cell types are coloured differently; visually, Figure 1(d) shows a better separation of cell types compared to Figure 1(c). This can be quantified by evaluating the accuracy of a nearest neighbour classifier on these two-dimensional data sets, as proposed in [21] (see Methods). The accuracy achieved by the classifier is reported inset in the individual panels in Fig 1(c-d), showing a stronger performance of CRAK-Velo in all the cases. Figure 1(e) shows histograms of the accuracy of such a classifier on all genes for the different methods. Methods using chromatin information retrieve in general a higher accuracy, but while this effect is modest for MultiVelo, CRAK-Velo provides a much greater accuracy for a very large number of genes.

Figure 1(f) depicts graphically the dynamics of regulation for two key genes, the transcription factor erythroid Kruppel-like factor 1 (KLF1) and the proteinase PRTN3, in terms of their proximal regulatory regions (within *±* 5kb from transcription start site). The plots show in blue the average unspliced read counts as a function of pseudo-time, while in other colours the effect of individual chromatin regions is displayed (obtained as the product of the open chromatin probability times the weight inferred by CRAK-Velo). In the case of KLF1, for example, it is evident that the increasing trajectory is driven by the high activity of a promoter-proximal region (orange), which then decreases sharply causing the observed transient behaviour. Next to the plots, a static picture showing the weight associated with each regulatory region and its position along the genomic region near the gene is shown as an arrow plot (as in e.g. [21]).

Next, we applied CRAK-Velo to another challenging data set, detailing mouse embryonic brain development at day 18 using 10X Multiome technology. While all three methods obtain broadly similar flows (Figure 2(a)), a detailed observation reveals subtle but relevant differences, particularly regarding the dynamics at the boundary between the terminally differentiated states of the Upper and Deeper Layer. We use the PAGA graph approach[22] to summarize transitions in Figure 2(b). Both MultiVelo and UniTVelo recover a spurious flow from Upper Layer to Deeper Layers, while CRAK-Velo correctly recognises these cell states as independent terminally differentiated states.This is appreciated more clearly in Figure 2(c), where the inset shows in detail the bifurcating flow at the boundary of subplate and Upper and Deeper Layer. An analysis of the inferred latent time (Figure 2a) also confirms this observation, as both MultiVelo and UniTVelo predict the deeper layers to develop after the upper layers, while CRAK-Velo assigns them both a similar pseudotime, consistent with biological knowledge. Limitations nevertheless remain, as all three methods were not able to identify ependymal cells as a terminal state. Next, we quantified the ability of the various methods to deconvolve cell types based on their inner representation. Figure 2(d-e) shows that chromatin-based approaches are more effective at detecting different cell types on three example genes, while Figure 2(f) quantifies across all genes the ability to separate cell types, once again confirming a significant superiority in performance for CRAK-Velo. Figure 2(g) finally depicts the regulatory dynamics of the Musashi RNA-binding protein 2 gene (MSI2). It is involved in stem cell development and is expressed in several structures, including the central nervous system. The figure again depicts a non-trivial regulatory process, providing a quantitative basis for understanding the behaviour of a key gene with a complex regulatory architecture.

**Fig. 2.**
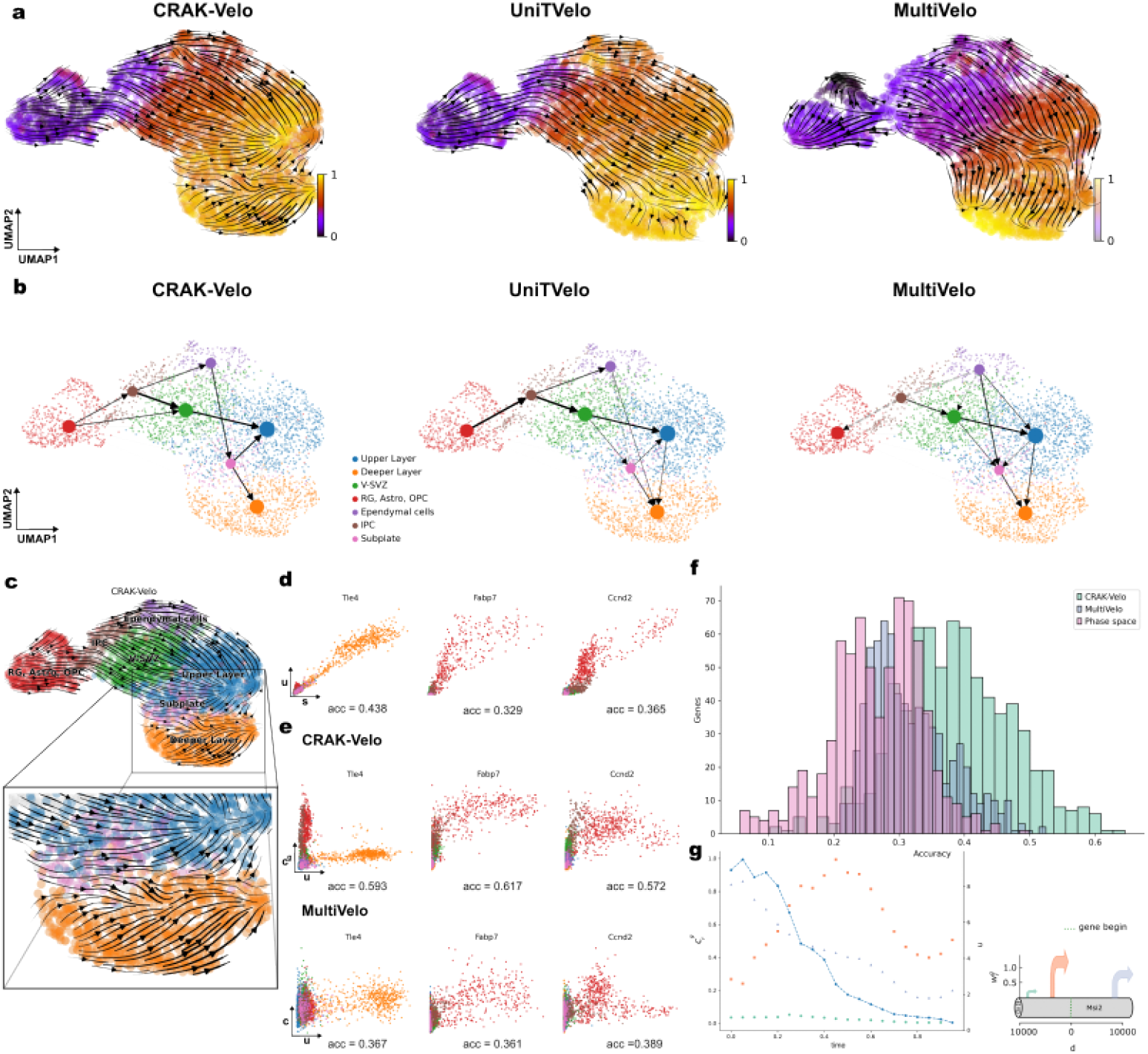
(a)UMAP visualisation of the velocity fields (shown in streamlines) inferred by CRAK-Velo, MultiVelo, and UniTVelo. The cells are colored according to the inferred pseudotime by the corresponding method. (b) PAGA graph summarizing transitions between different cell types obtained from the velocity field embeddings. (c) Detailed view of the bifurcating flow at the boundary of the subplate and Upper and Deeper Layers. (d-e) Three genes example(TLe4, Fabp7, and Ccnd2) to visualise cell type deconvolution based on un/spliced phase space in the upper row and chromatin accessibility-unspliced reads in the middle and bottom rows. The number below each plot is the accuracy of a KNN cell type classifier. (f) Histogram of the performance of the KNN cell type classifier across all genes for the phase space and accessibility-unspliced space inferred by the CRAK-Velo and MultiVelo. (g) Regulatory dynamics of the MSI2 gene. Left is the region kinetic plot with double y axis that shows the average accessibility 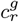 for each region around MSI2 on the left and the average normalised unspliced reads *u*_*g*_ on the right one. On the right is an arrow plot. The position of an arrow corresponds for start of a region and thickness is its contribution/weight 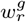 inferred by CRAK-Velo.

In summary, results on these two benchmark data sets show that CRAK-Velo is a promising tool to extract dynamic information from high-throughput single-cell multiomic data. It achieves accurate reconstruction of complex dynamic flows, and superior capabilities in cell-type deconvolution compared to the recent state-of-the-art method MultiVelo. Additionally, the simplicity of the method makes it straightforward to interpret the impact of changes in chromatin accessibility on transcriptional dynamics, generating potentially testable hypotheses on the regulation of gene expression in specific genes. CRAK-Velo therefore can extend the scope for biological insights deriving from the rapidly expanding field of single-cell multi-omics.

## 2 Methods

### 2.1 Data preprocessing

For the HSPC dataset, same cells and their corresponding annotations in MultiVelo are used. For the embryonic E18 mouse brain we followed the cell annotations of MultiVelo. All cells that belongs to the cell types list chosen by MultiVelo (‘RG, Astro, OPC’,’IPC’, ‘V-SVZ’, ‘Upper Layer’, ‘Deeper Layer’, ‘Ependymal cells’, ‘Subplate’) are kept for downstream analysis.

All the scRNA-seq part of the multi-omics datasets are preprocessed using the same approach as UniTVelo: The highly variable genes (HVGs) which contribute to cell-cell variation were selected (default 2,000). further selected informative genes with similar settings in scVelo.

For the scATAC-seq data part we chose to keep regions that are captured in more than 800 cells in the HSPC dataset and 400 cells for the embryonic E18 mouse brain dataset. Then regions that are within 10^4^ upstream and downstream from the gene starting site are kept.

The final HSPC dataset is composed of 11605 cells, 2000 genes, and 3939 regions. For the embryonic E18 mouse brain dataset we ended it up with 3365 cells, 2000 gene, and 4002 regions.

#### Smoothing scATAC-seq data

CisTopic is used to smooth the binary and sparse scATAC-seq data. We implemented our version of cisTopic in python (the link to our cisTopic implementation provided in the code availability section). CisTopic extracts a latent representation of dimension T for each cell n and region r:

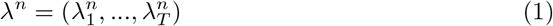

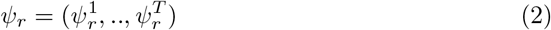

We computed the open probability of a region r in a cell n 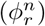 using the cisTopic representation:

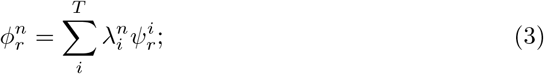

CisTopic is implemented with a Gibbs sampler. The number of samples is 3000 with thinning every 10 samples to avoid sample correlations. The extracted latent space has a dimension T=30 and the sample mean was used to estimate *λ*^*n*^ and *ψ*_*r*_,

### 2.2 Reconciling data-driven and chromatin-based transcription rates in velocity estimation

UniTVelo is based on a data-driven, parametric estimate of the amount of spliced reads as a function of time (using flexible basis function regression). Following the default workflow of UniTVelo, the spliced read of a gene is modelled explicitly using RBF basis function regression as [12]:

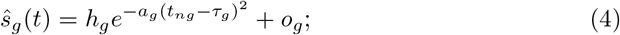

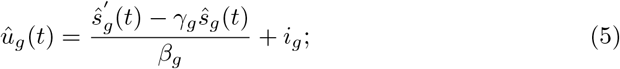

Applying time derivative on equation 5, we obtain 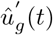 written in terms of *Ŝ*_*g*_(*t*) and its time derivative 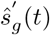:

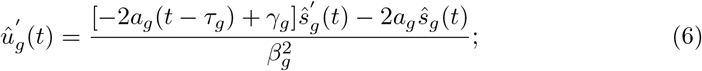

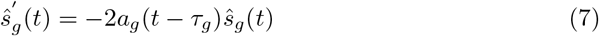

Alternatively, we can use scATAC-seq to define the production rate of unspliced reads as follows. The scATAc-seq data represents the accessibility information of the cis-regulatory regions across the genome. Fixing a window around a specific gene *g* we obtain a set of regions of size *R*_*g*_. Using this set of regions, we define the ATAC-derived transcription rate of gene *g c*^*g*^ as follows:

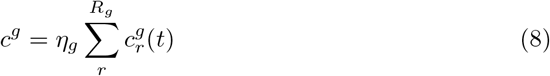

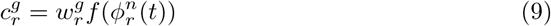

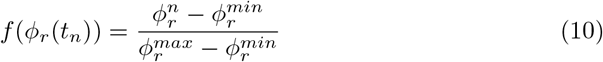

Time *t*_*n*_ is the inferred unified pseudotime for a cell *n*. The dynamic variable 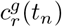 represents the accessibility of a region *r* associated with a gene *g* in cell *n* which has a pseudotime *t*_*n*_. Periodically the accessibility function is constructed 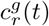 by reordering the cell’s accessibility according to the inferred pseudotime of the cells (more details in the parameter inference section). Different regions around a specific gene contribute with different weights to the regulation of this gene. A region *r* around gene *g* regulates this gene with a weight 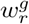 that need to be inferred. In all the results shown *R*_*g*_ = 10^4^bp.

Therefore, we can define the ATAC-derived dynamics of unspliced read counts as

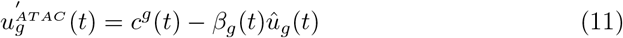

We end up with two forms of the derivative of unspliced reads. The unspliced derivative that is derived from the proposed paramtrised RBF (equation 6) while the other form encodes the scATAC-seq data information within it (equation 11). In the next section, we show how to reconcile it through a bespoke likelihood function.

### 2.3 Parameter inference of CRAK-Velo

In order to reveal the accessible chromatin region and splicing kinetics we need to infer gene-specific parameters *θ*_*g*_ and cell-specific time points *t*_*ng*_. The set of paramters, *θ*_*g*_, are used to define the shape of mean function of both un/spliced reads, where *θ*_*g*_ is:

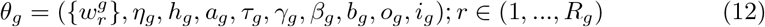

Let 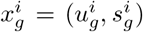 where 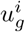 and 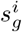 are the normalized observation of un/spliced counts for a particular gene and *i* ∈ (1, …, *N*) such that *N* is the number of cells. Similarly, let 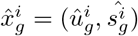 be the model’s estimation of un/spliced counts of that gene.

The objective function used to find the optimal parameter settings of phase portrait and the accessible region kinetics is negative log-likelihood that includes multi-omics information of the un/spliced reads and scATAC-seq smoothed data:

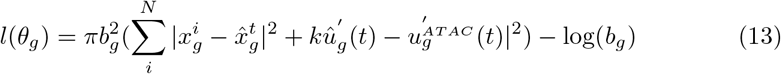

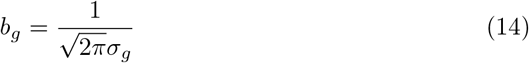

where k = 0.5 to tune the effect of noise coming from the scATAC-seq data on the inference procedure, and *σ*_*g*_ is defined in UniTVelo as the standard deviation of the Euclidean distance residuals under the assumptions that the residuals are normally distributed 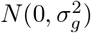.

We used the same optimization procedure in UniTVelo[12] to fit the model parameters by iteratively minimizing the negative log-likelihood using Gradient Descent algorithm:

- Given 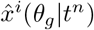, parameterized by unified cell time, the gradients of the objective function is computed and *θ*_*g*_ is updated.
- For a fixed *θ*_*g*_ gene-cell specific time *t*_*ng*_is reassigned by minimizing the Eculidean distance between *x*^*i*^ and updated 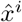.
- The unified pseudotime of the cells *t*_*n*_ is computed.
- The variable *ϕ*_*r*_ for a region r is ordered according to the unified pseudotime. Note that the loss of the genes that has no regions around within the chosen window is the same as the UniTVelo one.

We used non-informative initialization as defined in UniTVelo with initial value *t*_*n*_ = 0.5.We run for 10000 epochs for each dataset.

### 2.4 Evaluating model performance

We used cross boundary direction on the HSPC dataset to evaluate the performance of CRAK-Velo, UniTVelo, and Multivelo in capturing meaningful differentiation trajectories of the hematopoietic stem cells.

We used K-nearest neighbors(KNN) classifier to evaluate CRAK-Velo and MultiVelo performances in deconvoluting cell-types once the inferred accessibility information c is used. Also we compare it with the observations (un/spliced reads) to quantify the improvement the accessibility can add to the observed data. The number of neighbors used in KNN is 5 and the metric is Euclidean. We trained the KNN on 70% of the cells in each dataset and validated on the rest. We show in the histograms the genes that are common between CRAK-Velo and MultiVelo.

The PAGA graph in Figure 2b is obtained by using latent time as a time prior and velocity field embeddings inferred by the three methods. All the possible edges obtained through PAGA by setting threshold = 0 and minimum spanning tree option to False.

### 2.5 Region kinetics

We monitored the accessibility kinetics of regions around a chosen genes by dividing the timeline (between 0 and 1) into intervals of width 0.05 and compute the average accessibility 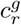 per interval. We showed in parallel the change in the unspliced reads with cell pseudotime by also average the normalised unspliced reads over an interval of 0,05. The kinetic plots have a double y axis one that shows the accessibility on the left and the normalised *u*_*g*_ on the right one.

### 2.6 Data availability

10x embryonic mouse brain dataset can be accessed at the 10x website at https://www.10xgenomics.com/resources/datasets/fresh-embryonic-e-18-mousebrain-5-k-1-standard-1-0-0. https://www.10xgenomics.com/resources/datasets/fresh-embryonic-e-18-mouse-brain-5-k-1-standard-1-0-0.

The processed files of the newly sequenced 10x Multiome HSPC samples are available at the GEO (GSE209878). The instructions to download the preprocessed version of the data by MultiVelo utilized in our analysis is available in the Multivelo tutorials.

### 2.7 Code availability

CRAK-Velo code is freely available on github https://github.com/StatBiomed/CRAK-Velo.git. Detailed workflows to reproduce figures and results in this paper are written as Jupyter notebook in the repository. Our implementation of cisTopic is available here: https://github.com/Nour899/cisTopic.git.

## Acknowledgements

We thank Fabian Theis and Weixu Wang for useful discussions towards the beginning of this project, and Maria Ayub for help in interpreting the biological results.

